# Cytokinins specify pluripotent stem cell identity in the moss *Physcomitrium patens*

**DOI:** 10.1101/2023.12.19.572286

**Authors:** Yuki Hata, Juri Ohtsuka, Yuji Hiwatashi, Satoshi Naramoto, Junko Kyozuka

## Abstract

The shoot apical meristem (SAM) contains pluripotent stem cells that produce all the aerial parts of the plant. Stem cells undergo asymmetric cell divisions to self-renew and to produce differentiating cells. Our research focused on unraveling the mechanisms governing the specification of these two distinct cell fates following the stem cell division. For this purpose, we used the model organism *Physcomitrium patens*, which features a singular pluripotent stem cell known as the gametophore apical cell. We show that the activity of cytokinins, critical stem cell regulators, is restricted to the gametophore apical cell due to the specific localization of PpLOG, the enzyme responsible of cytokinin activation. In turn, PpTAW, which promotes differentiating cell identity of the merophyte, is excluded from the gametophore apical cell by the action of cytokinins. We propose a cytokinin-based model for the establishment of asymmetry in the pluripotent stem cell division.

**One Sentence Summary:** Cytokinins are confined to the pluripotent stem cells and exclude the differentiation factor TAW1 to establish the SAM from a single stem cell.

## Main Text

All the diverse tissues and organs in multicellular organisms originate from stem cells (1–3). In plants, stem cells in the shoot apical meristem (SAM) underlie aerial shoot growth (4). The root apical meristem is the originator of all of the cells in the root system (5). The advent of stem cells that can produce distinct cell types has been the driving force behind the remarkable morphological diversity of plants, which enabled the colonization of terrestrial environments and the success of land plants (6–9). In flowering plants (angiosperms), a group of stem cells is maintained at the center of the SAM, while the SAM of bryophytes and most ferns contain a single stem cell known as the apical cell (Fig. 1A) (10,11). Studies have shown that core regulatory mechanisms controlling the function of the SAM are conserved in land plants in spite of the differences in SAM structure (12–20). Cytokinins, a class of phytohormones, has been shown to be a common factor promoting stem cell identity within the SAM in land plants (19). A crucial role of the stem cell is to perpetuate asymmetric cell divisions to ensure both self-renewal and the production a progenitor cell that will differentiate (3). However, the precise mechanism by which cytokinins specify the fate of the stem cells and control the asymmetric division of these cells in plants remain largely unknown.

**Fig. 1.**
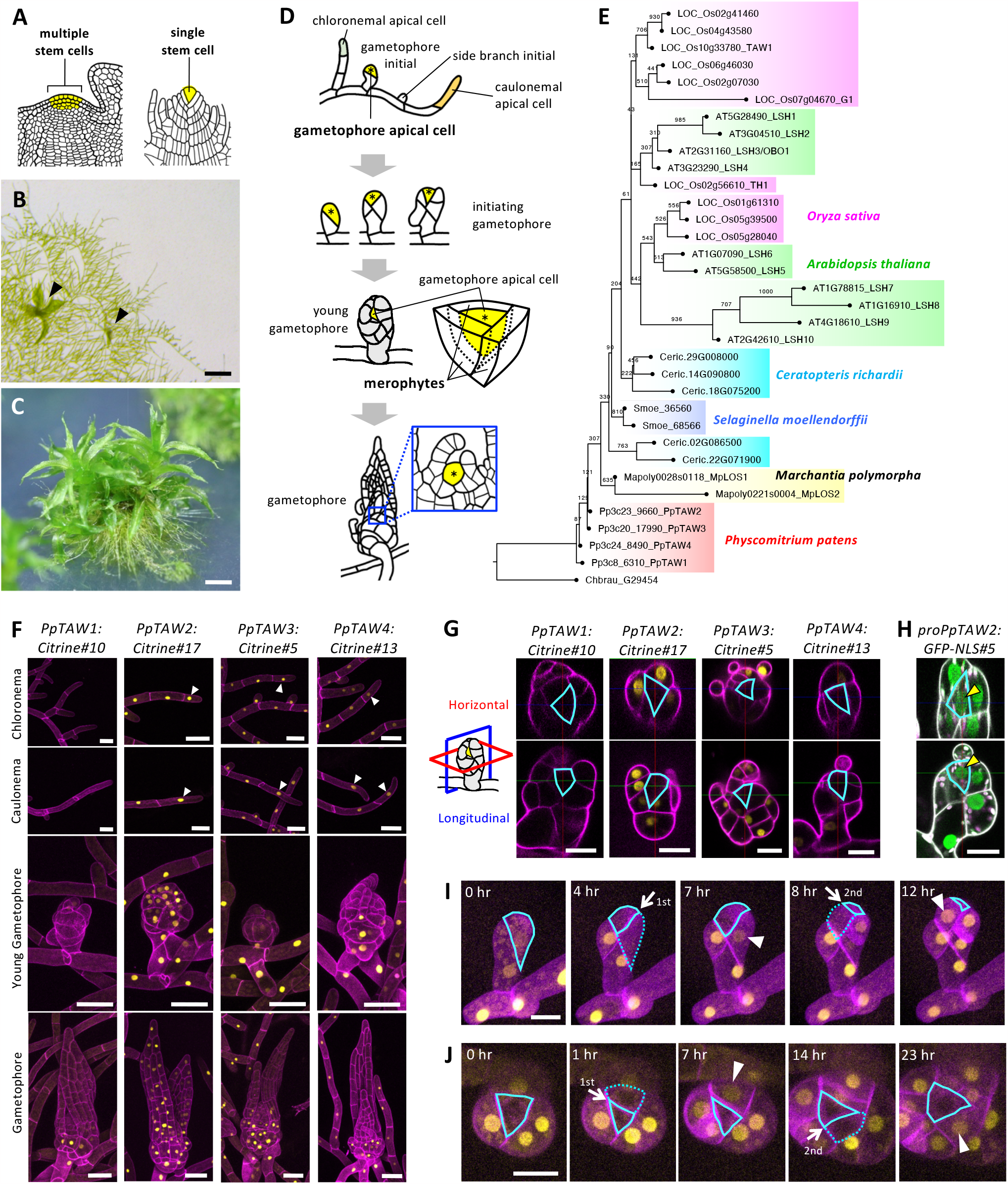
The PpTAW protein is excluded from the gametophore apical cell through post-transcriptional regulation. (**A**) Structure of shoot apical meristem (SAM) in land plants. Stem cells are indicated by yellow color. (**B** and **C**) Growing protonemata of *Physcomitrium patens*, possessing young gametophores (B) and mature gametophores (C). Black arrowheads indicate young gametophores. (**D**) Schematic representation of gametophore initiation and development. Some of the side branch initials are specified as the gametophore apical cell and grow into the gametophore initial. After a defined pattern of cell divisions, the gametophore initial grows into the gametophore. The gametophore apical cell resides at the center, surrounded by merophytes, which will become the leaves. Yellow cells with an asterisk indicate the gametophore apical cell. (**E**) Phylogenetic tree of ALOG proteins in land plants. Species shown in the tree are *Chara braunii* (Chbrau), *Physcomitrium patens* (Pp), *Marchantia polymorpha* (Mapoly), *Selaginella moellendorffii* (Sm), Ceratopteris richardii (Ceric), *Arabidopsis thaliana* (AT), and *Oryza sativa* (Os). An ALOG protein of *Chara braunii* (Chbrau_G29454) was included as an outgroup. The bootstrap value is indicated at the branch point. (**F)** Localizations of PpTAW:Citrine fluorescence (yellow) in protonema, young gametophore and gametophore. PpTAW:Citrine fluorescence is visible in the chloronemal and caulonemal apical cells (white arrowheads). Cell walls were stained with propidium iodide (magenta). (**G**) Localizations of PpTAW:Citrine fluorescence (yellow) in the shoot apical meristem (SAM). The top panels show horizontal (red square) views, and the bottom panels show longitudinal (blue square) views. Cell walls are stained with propidium iodide (magenta). The outlines of the gametophore apical cell is marked with cyan lines. (**H**) Promoter activity of *PpTAW2*. GFP fluorescence (green) driven by the *PpTAW2* promoter is shown. Cell walls are stained with propidium iodide (white). The magenta color represents the autofluorescence of chloroplasts. Yellow arrowheads indicate the GFP fluorescence in the gametophore apical cell. (**I** and **J**) Time-lapse imaging of PpTAW2:Citrine (yellow) localization during the division of gametophore apical cells in an initiating gametophore (I) and a growing young gametophore (J). LTI6b:RFP (magenta) was simultaneously imaged for the visualization of cell outlines. Solid and dashed cyan lines outline the gametophore apical cell and newly formed merophyte, respectively. White arrows indicate the cell division planes formed during the observation time. White arrowheads indicate the PpTAW2:Citrine signal that appeared in the merophyte after division of the gametophore apical cell. No PpTAW2:Citrine signal is observed in the gametophore apical cell. Scale bars, 500 μm (B), 300 μm (C), 50 μm (F), and 20 μm (G to J).

*Physcomitrium patens* is a suitable model for studying SAM development (Fig. 1, B and C) (21,22). The SAM of *P. patens* exhibits a simple structure with clear cell types and is applicable to live imaging. The cell division pattern in the SAM, which has been well documented, is highly coordinated (23). During the life cycle of *P. patens*, filaments called chloronemata grow following the germination of spores. The chloronemal apical cell then undergoes a transition to become the caulonemal apical cell. Chloronemata and caulonemata, are collectively called protonemata. The growth of the protonemata is characterized by the division and extension of the chloronemal and caulonemal apical cells at their tips (Fig. 1D). Most side branch initials generated on the protonema grow as filaments, contributing to the two-dimensional expansion of the plant. Nevertheless, a subset of these side branch initials gives rise to gametophores (Fig. 1D, gametophore initial). The gametophore apical cell is a pluripotent stem cell, different from protonemal apical cells, which possess limited stem cell capability for self-renewal (21). The gametophore apical cell follows a characteristic division pattern, resulting in the formation of a SAM featuring a central tetrahedral apical cell and an outer cell known as the merophyte. The merophyte differentiates to form a leaf and a segment of the stem (Fig. 1D, gametophore).

### PpTAW proteins are excluded from the gametophore apical cell

ALOG (Arabidopsis light-dependent short hypocotyl (LSH) and *Oryza* G1) proteins are transcription factors conserved in land plants (24). *ALOG* genes exhibit high sequence similarity across the coding region among land plants (fig. S1). In angiosperms, *ALOG* genes are expressed in the boundary region that lies between the SAM and differentiating organ primordia. They regulate SAM indeterminacy, boundary differentiation, and lateral organ size (25–29). In the bryophyte *Marchanthia polymorpha*, a single ALOG gene, *LATERAL ORGAN SUPRESSOR 1* (Mp*LOS1*), directly controls the identity of differentiating cells and indirectly the activity of the gametophyte apical cell (30).

There are four *ALOG* genes in the *P. patens* genome, namely, *PpTAW1* (Pp3c8_6310), *PpTAW2* (Pp3c23_9660), *PpTAW3* (Pp3c20_17990), and *PpTAW4* (Pp3c24_8490) (Fig. 1E and fig. S1). We first examined the localization of the four PpTAW proteins using reporter lines (*PpTAW1:Citrine#10, #12, PpTAW2:Citrine#17, #9, PpTAW3:Citrine#5, #2, PpTAW4:Citrine#13, #15*) in which the Citrine gene is inserted at the C-terminus of the coding sequence. Consistent with the function of ALOG proteins as transcription factors, the fluorescence of the fusion protein localized in the nucleus in all lines. Although the signal intensity varied among the four *PpTAW*s; with the highest observed for the *PpTAW2:Citrine* fusion and the lowest for the *PpTAW1:Citrine* fusion, all *PpTAW:Citrine* reporter lines showed fluorescence in almost all tissues, including the protonemata, side branch initials, and gametophores (Fig. 1F and fig. S2A). We further examined the localization of the PpTAW:Citrine signal within the gametophore apical cell in the SAM. The gametophore apical cell is shaped as a regular tetrahedron, thus it appears as a triangular shape at the center of the SAM in both the horizontal and longitudinal sections. The signal of all four PpTAW:citrine fusion proteins was excluded from the gametophore apical cell (Fig. 1G and Fig. S2B). Since we confirmed that the spatial localization patterns of all four PpTAW proteins are similar, we focused our subsequent analysis on *PpTAW2:Citrine*, which showed the strongest signal. To test if the PpTAW2 protein localization is regulated at the transcriptional level, we generated *proPpTAW2:GFP-NLS* lines expressing nuclear-localized GFP under the control of the *PpTAW2* promoter. The GFP fluorescence was detected in the gametophore apical cell, indicating that the transcription of *PpTAW2* occurs in the gametophore apical cell (Fig. 1H, and fig. S2, C and F). These results suggest that post-transcriptional regulation of *PpTAW2* expression takes place, inhibiting PpTAW2 protein accumulation in the gametophore apical cell. In contrast, all PpTAW:Citrine fusion proteins were found to accumulate in the protonemal apical cells (Fig. 1F and fig. S2A). These observations suggest a correlation between the absence of the PpTAW2 protein and gametophore apical cell identity.

We performed live imaging analysis of the PpTAW2:Citrine fusion protein in the gametophore apical cell during the early stages of gametophore initiation (Fig. 1I, fig. S2D, movie S1, and movie S2). The PpTAW2:Citrine signal was absent in the gametophore apical cell, while a weak signal was detected in the merophyte at the two-cell stage (0 hr). After the division of the gametophore apical cell (4 hr in Fig. 1I), the PpTAW2:Citrine signal became visible in the merophyte cell but not in the apical cell (7 hr in Fig. 1I). This pattern was repeated in the following divisions of the gametophore apical cell. As the gametophore continued to grow, PpTAW2:Citrine fluorescence appeared only in the cell destined to become the merophyte following the division of the apical cell (Fig. 1J, fig. S2E, movie S3, and movie S4). These results indicate that the PpTAW2 protein was effectively removed or absent from the gametophore apical cell from the first division of the gametophore apical cell and the pattern of PpTAW2 localization persisted throughout gametophore development.

### Cytokinins repress PpTAW2 protein accumulation in the gametophore apical cell

The PpTAW2 protein is present in the protonemal apical cells, which are uni-potent stem cells, but is absent in the gametophore apical cell, a pluripotent stem cell. Cytokinins promote the generation of the gametophore apical cells but do not induce the generation of protonemal stem cells (12,19,31,32). These prompted us to investigate whether PpTAW2 protein accumulation is suppressed by cytokinins in the gametophore apical cell. To explore this possibility, we first treated plants expressing the PpTAW2:Citrine fusion with a cytokinin (6-benzylaminopurine (BAP)) and subsequently confirmed the induction of *PpCKX1*, an ortholog of *CYTOKININ OXIDASE 1* (*CKX1*), a cytokinin-responsive gene (19). The intensity of the PpTAW2:Citrine fluorescence was significantly decreased in young gametophores treated with BAP (Fig. 2, A and B) while the endogenous *PpTAW2* mRNA levels were not affected (Fig. 2C). These results support our assumption that PpTAW2 protein accumulation is repressed posttranscriptionally by cytokinins. There are several possible mechanisms for post-transcriptional repression of PpTAW2. These mechanisms may include inhibition of translation, protein degradation, and symplastic transportation from the gametophore apical cell. Investigating these mechanisms should be the next step in gaining a better understanding of how PpTAW2 expression is regulated.

**Fig. 2.**
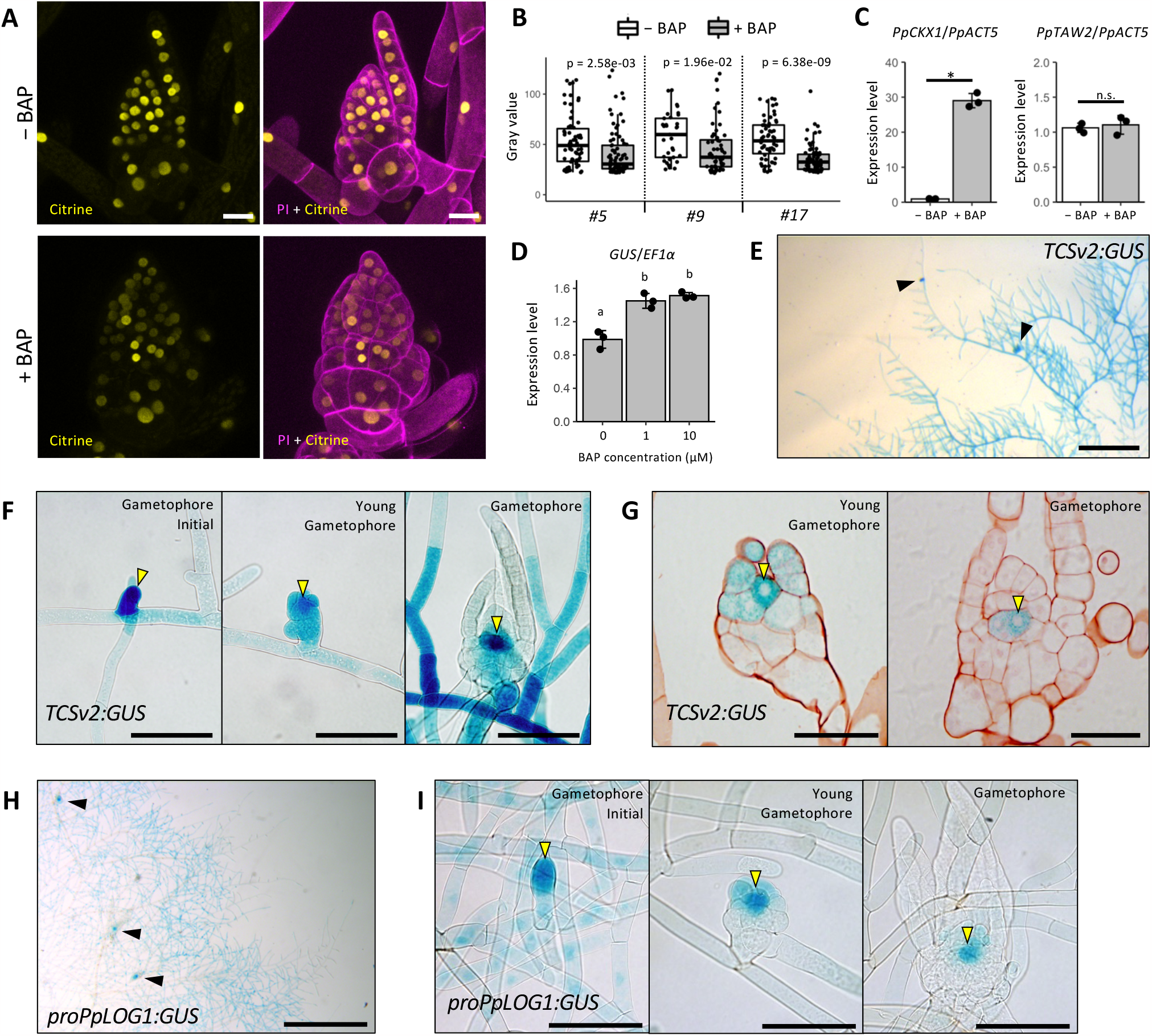
Cytokinins localize in the gametophore apical cell and repress PpTAW2 protein accumulation. (**A**) Effects of cytokinin (BAP) treatment on the accumulation of PpTAW2:Citrine (yellow). The outline of cells is visualized with propidium iodide (magenta). (**B**) Signal intensity of PpTAW2:Citrine in control (−BAP) or cytokinin-treated (+ BAP) gametophores in three independent lines. Citrine fluorescence in nuclei was measured by quanifying using imageJ. Three gametophores were used in each condition. Statistical significance was examined using Student’s t-test (n ≧ 33). (**C**) Effects of cytokinin (BAP) treatment on the *PpTAW2* mRNA expression. mRNA levels of *PpCKX1* and *PpTAW2* were quantified by qPCR. Statistical significance was assessed using Student’s t-test (n = 3, p < 0.05). (**D**) Response of *TCSv2:GUS* expression to the cytokinin (BAP) measured by qRT-PCR. Statistical significance was evaluated by HSD test (n = 3, p < 0.05). (**E-G**) GUS activity of *TCSv2:GUS* in whole tissues (E), developing gametophores (F), and transversal sections of SAM in the gametophores (G). Black arrowheads in (E) indicate gametophores formed on protonema. Yellow arrowheads in (F) and (G) indicate the gametophore apical cell. (**H-I**) GUS activity of *proPpLOG1:GUS* in whole tissues (H) and developing gametophores (I). Black arrowheads indicate gametophores formed on protonemata. Yellow arrowheads indicate strong GUS signal observed at the gametophore apical cell. Scale bars, 20 μm (A), 50 μm (G), 100 μm (F and I), and 500 μm (E and H).

We examined the spatial localization of cytokinins in the SAM using the *TWO COMPONENT SENSOR version2* (*TCS2*) system with GUS (β-glucuronidase) as a reporter (33). First, we confirmed the responsiveness of GUS expression to BAP treatment (Fig. 2D). The *TCSv2:GUS* lines showed a weak to moderate GUS signal in most protonemal cells. In contrast, intense signals were observed in the gametophore apical cells and gametophores at the early stage (Fig. 2, E and F). As the gametophore grew, the GUS signal became restricted to its central part (Fig. 2F). We further confirmed the presence of strong GUS expression in the gametophore apical cell in the young and growing gametophore (Fig. 2G). These analyses revealed that intense cytokinin signaling occurred in the gametophore apical cell from the singlecell stage, and the cytokinin signaling was consistently maintained in the gametophore apical cell as it continued to develop.

LONELY GUY (LOG) catalyzes the final step of the cytokinin biosynthesis pathway (34). *P. patens* possesses nine genes encoding LOG orthologs (Fig. S3) (35). To clarify the localization of cytokinin biosynthesis, we generated marker lines in which the GUS gene is fused with the sequence of the *PpLOG1* gene promoter, which exhibits a strong activity (40, 41). Strong GUS activity was detected in the gametophore initial cell and the gametophore apical cell in the SAM, similar to the GUS signal localization observed with the *TCSv2:GUS* lines (Fig. 2, H and I). These results suggest that specific localization of *LOG* expression in the gametophore apical cell restricts cytokinin activity to the gametophore apical cell. The simultaneous localization of cytokinin signaling and PpTAW2 protein and the suppression of PpTAW2 protein accumulation by cytokinins provides strong support to our hypothesis that PpTAW2 protein accumulation is inhibited by cytokinins in the gametophore apical cell.

### Inhibition of *PpTAW*s’ function causes overgrowth of protonemata and leaves and impairs merophyte differentiation

To reveal the function of *PpTAW*s, we generated loss-of-function mutants by homologous recombination. Double mutants of *PpTAW2* and *PpTAW3* (*Δpptaw2Δpptaw3*) showed a significant increase in colony size. The increase in colony size was intensified in lines bearing mutations in multiple *PpTAW* genes and it reached its highest level in the *quadruple* mutants (Fig. 3, A to C, and fig. S4A). The defect was caused by an increase in both the number of cell divisions and the length of the protonemal cells. Colony growth in *P. patens* exclusively depends on the tip growth and division of protonemal apical cells (21). Thus, PpTAWs inhibit the activity of protonemal apical cells to elongate and divide.

**Fig. 3.**
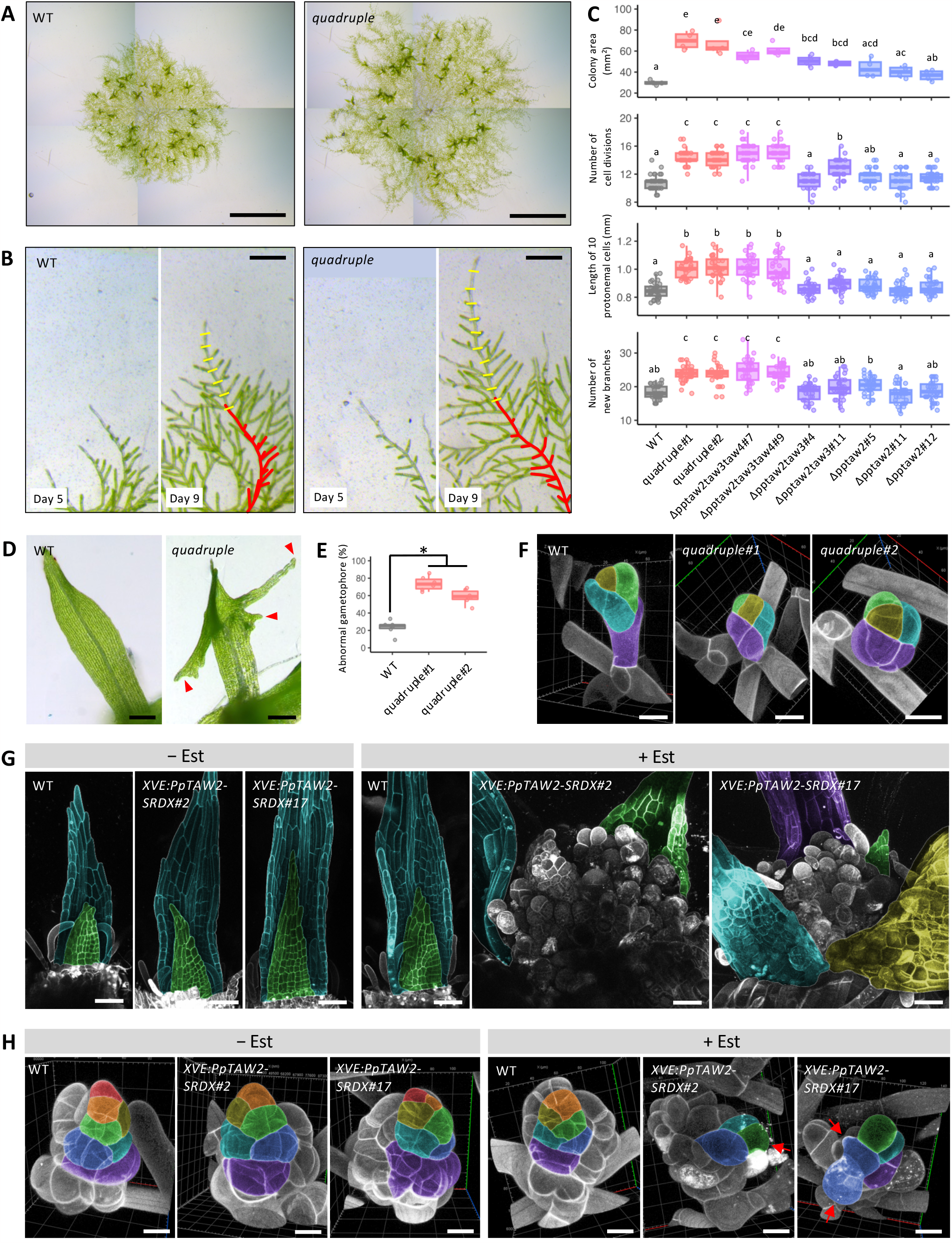
Disruption of *PpTAW*s’ function causes overgrowth of protonemata and abnormal organ differentiation. (**A**) Colonies of WT and *PpTAW* quadruple mutant at 23 days of culture. (**B**) Growth of protonemata of WT and the quadruple mutant from day 5 to day 9 of culture after inoculation. Red lines on the day 9 plants outline the silhouettes of day 5 plants. Yellow lines on the day 9 plants indicate the position of newly formed cell wall by division of the caulonemal apical cell. (**C**) Quantification of colony growth of *PpTAW* loss-of-function mutants. Colony area of day 23 plants (n = 4), number of caulonemal apical cell divisions between day 5 and day 9 after inoculation (n = 30), length of caulonemal cells (n = 30), and number of new branches formed on a caulonemal filament (n = 30) in WT and *PpTAW*s loss-of-function mutants. Statistical significance was evaluated using the HSD test (p < 0.05). (**D**) Leaves in gametophores of WT and quadruple mutants. Red arrowheads indicate abnormal elongation of leaf margins. (**E**) Frequency of gametophores showing abnormal leaf elongation in WT and quadruple mutants. Statistical significance was examined by student’s t-test (n = 6, p < 0.05). (**F**) Confocal images of initiating gametophores in WT and quadruple mutants. The gametophore apical cell is shown in yellow. Green, magenta, and cyan-colored cells are progenitor cells derived from different merophytes. (**G**) Effects of PpTAW2-SRDX induction of gametophore development. Confocal images of the gametophore apex in WT and *XVE:PpTAW2-SRDX* lines are shown at 4 weeks after β-estradiol application (0 or 100 nM). Green, cyan, magenta, and yellow-colored tissues are leaf primordia or young leaves. (**H**) Cell division patterns of merophytes in WT and *XVE:PpTAW2-SRDX* lines at 4 weeks after application of 100 nM β-estradiol (+ Est) or solvent (− Est). Clonal sectors derived from the same merophyte are shown using the same color. Red arrows indicate irregular cell elongation. Cell walls are visualized by propidium iodide staining in (F), (G), (H). Scale bars, 4 mm (A), 200 μm (B and D) 20 μm (F and H), and 50 μm (G).

Continuous observation of the same plants revealed that the number of protonemal branches generated between Day 5 and Day 9 was higher in the quadruple mutants than in WT (Fig. 3, B and C). These results indicate that the four *PpTAW* genes redundantly function in repressing division and growth of protonemal apical cells and generation of side branch initials.

Gametophore development was also affected in the quadruple mutants. Leaves, which originate from the merophyte, undergo coordinated cell divisions and growth to take a flat and elliptical shape (38). In the quadruple mutants, the leaves frequently exhibited excessive growth (Fig. 3, D and E). These observations suggest that *PpTAW*s play a crucial role in regulating the timing of cell divisions. The rhizoid, a filamentous tissue at the base of the gametophore, gradually becomes brown as it matures (39). The browning of rhizoid cells is inhibited in the quadruple mutants, indicating that the *PpTAW*s genes are also required to promote rhizoid cell maturation (Fig. S4, B and C).

We did not observe morphological alterations in the SAM of gametophores in the quadruple mutants (Fig. 3F). To further investigate the role of PpTAWs as transcription regulators in SAM development, we generated β-estradiol-inducible lines expressing *PpTAW2-SRDX*. PpTAW2-SRDX dominantly disturbs the normal function of PpTAWs to regulate downstream genes (40). In these lines, the expression of PpTAW2-SRDX is ubiquitously induced by β-estradiol application (41). Induction of *PpTAW2-SRDX* during gametophore development caused the proliferation of irregular cell masses at the top of the gametophore (Fig. 3G). Continuous observation of leaf development in these plants revealed that the growth pattern of cells in the leaf primordia was severely disturbed. However, the gametophore apical cell looked almost normal in *PpTAW2-SRDX* lines (Fig. 3H). ALOG proteins function both as transcriptional activators and repressors of downstream genes (42–44). If PpTAWs also act as both activators and repressors, disturbance of PpTAWs function in the SRDX system should result in pronounced disorders in gametophore development. However, we cannot rule out the possibility that abrupt inhibition of PpTAWs function caused the disturbance of gametophore development as it requires strict spatiotemporal regulation of gene expression. Further analysis of PpTAWs molecular function is necessary to distinguish between these two possibilities. In summary, the analysis of the quadruple mutants and *PpTAW2-SRDX* over-expression lines suggests that PpTAWs suppress the tip growth and division of the protonemal apical cells and formation of side branch initials while promoting the establishment of merophyte identity.

### Ectopic over-expression of PpTAW2 suppresses protonemal growth and stem cell initiation

We further analyzed the function of PpTAWs and the significance of their proper spatial localization using PpTAW2 inducible lines, *XVE:PpTAW2*, in which PpTAW2 can be induced by application of β-estradiol (41). We confirmed the presence of PpTAW2 in all tissues including the gametophore apical cell following β-estradiol treatment by using a *XVE:PpTAW2*-*Citrine* line (Fig. S5). Induction of XVE:PpTAW2 by β-estradiol application resulted in colony size reduction in a β-estradiol concentration dependent manner, suggesting an inhibitory effects of PpTAW2 on protonemal apical cell activity (Fig. 4, A and B). Differentiated tissues of *P. patens* retain the capacity to form new protonemal apical cells and to regenerate protonemata (45). We also examined the effects of PpTAW2 over-expression on protonemal apical cell regeneration. *XVE:PpTAW2* gametophores were cultured in the presence of β-estradiol, and then leaves were cut and cultured on the medium without β-estradiol. The formation of protonemal apical cells from the section of the cut leaves was suppressed in plants treated with β-estradiol in a concentration-dependent manner (Fig. 4, C and D). This provides further support for the inhibitory role of PpTAW2 in specifying protonemal stem cell identity and activity. In addition, applying a higher concentration (0.5 nM) of β-estradiol suppressed the formation of side branch initials (Fig. 4, E and F). Gametophore formation was almost fully suppressed by applying 1 nM β-estradiol, suggesting that PpTAWs inhibit the specification of gametophore apical cell identity and/or its activity (Fig. 4, A and G).

**Fig. 4.**
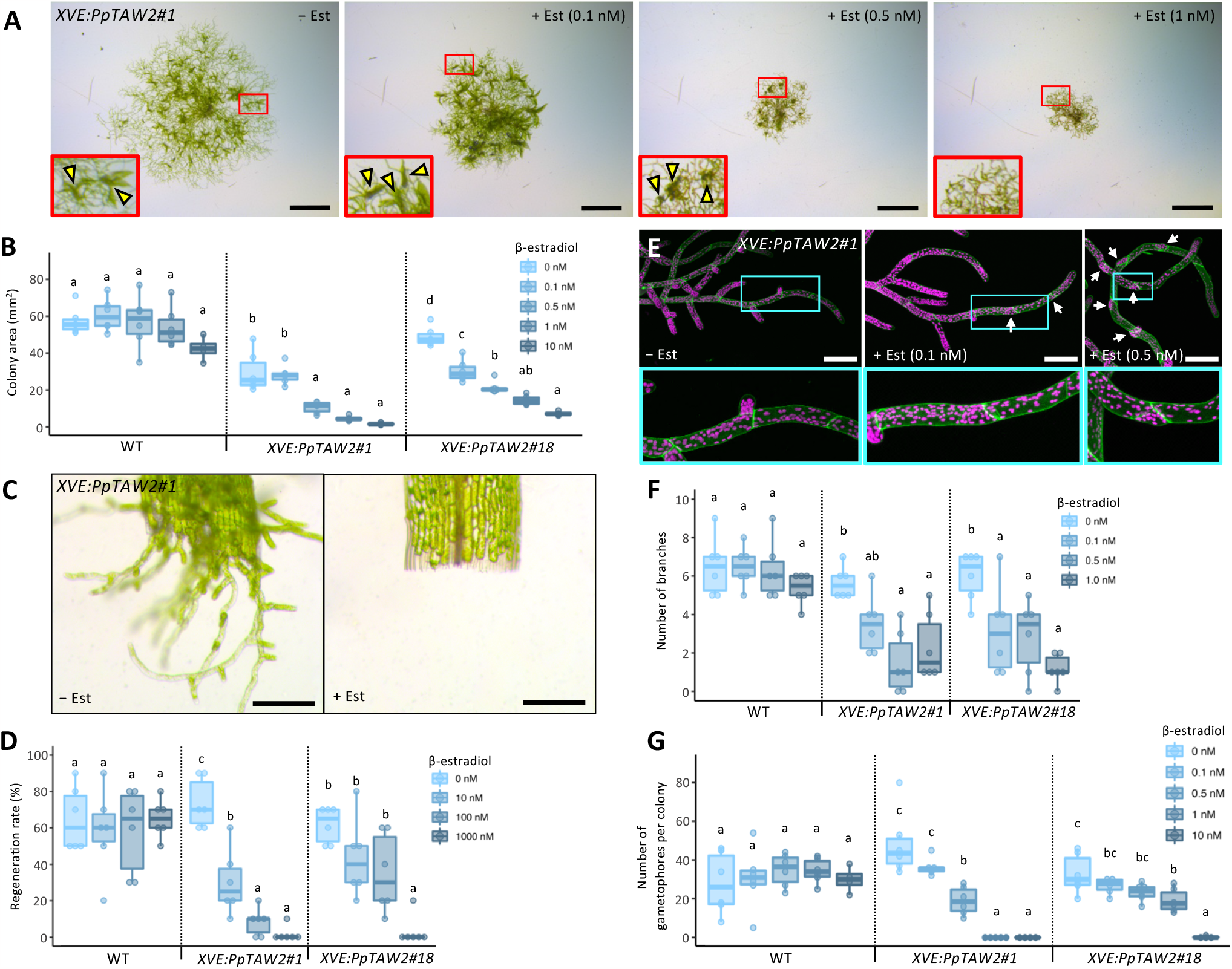
Ectopic over-expression of PpTAW2 inhibits protonemal apical cell activity and gametophore initiation. (**A**) Colonies of *XVE:PpTAW2* plants grown with (+ Est) or without (− Est) β-estradiol for 21 days. A close-up view of the framed area is shown at the lower-left corner. Yellow arrowheads indicate gametophores. (**B**) Colony area of WT and *XVE:PpTAW2* plants grown with different concentrations of β-estradiol for 21 days. Statistical significance was assessed using the HSD test (p < 0.05, n = 4-6). (**C**) Protonema regeneration from detached leaves of *XVE:PpTAW2* plants treated with 1μM β-estradiol (+ Est) or without β-estradiol (− Est), and grown for 3 days. (**D**) Frequency of protonemal regeneration from detached leaves in WT and *XVE:PpTAW2* lines3 days after treatment with different concentrations of β-estradiol. Statistical significance was assessed using the HSD test (p < 0.05, n = 6). (**E**) Confocal images of the protonemata of *XVE:PpTAW2* plants treated with (+ Est) or without (− Est) β-estradiol. The concentration of β-estradiol is shown in each upper subpanel. Closeup views of the cyan framed areas in the upper panels are shown in the bottom panels. White arrows indicate sites where the growth of side branch initials is arrested. Cell walls are visualized using propidium iodide (green). Magenta color is the autofluorescence of chloroplasts. (**F**) Quantification of the suppression of side branch initial elongation observed from the 9th caulonemal cells to the tip of the gametophore of WT and *XVE:PpTAW2* lines grown with different concentration of β-estradiol. Statistical significance was assessed using the HSD test (p < 0.05, n = 6). (**G**) Number of gametophores of WT and *XVE:PpTAW2* plants grown with different concentrations of β-estradiol for 21 days. Statistical significance was assessed using the HSD test (p < 0.05, n = 4-6). Scale bars, 2 mm (A), 200 μm (C), and 100 μm (E).

## Discussion

We showed that cytokinin activity is localized to the gametophore apical cell in the SAM of *P. patens*. Conversely, PpTAW promotes merophyte identity and is excluded from the gametophore apical cell due to cytokinin activity. Based on these findings, we propose a model to describe how the SAM of *P. patens*, which contains a single pluripotent stem cell known as the gametophore apical cell, is initiated and maintained (Fig. 5). Cytokinins work as pluripotent stem cell factors while PpTAW is a differentiation factor required to specify the merophyte identity. In addition, cytokinins suppress the accumulation of PpTAW proteins. Cytokinin activity is restricted to the gametophore apical cell as consequence of the localization of LOG, the final enzyme of the cytokinin biosynthesis pathway, within this cell, resulting in PpTAW being specifically excluded from the gametophore apical cell. This process leads to the autonomous establishment of asymmetric cell identity, pluripotent stem cell activity of the gametophore apical cell, and differentiating cell activity of the merophyte following the division of the gametophore apical cell. In this system, the key factor for establishing asymmetry is the confinement of cytokinin activity to the gametophore apical cell, which is accomplished by the specific expression of PpLOG. The next critical step in understanding the nature of the pluripotent stem cell is the elucidation of the mechanisms underlying the specific localization of *PpLOG. LOG* genes are expressed in the SAM, which contains the stem cell zone of angiosperms (34,46). Our finding that *PpLOG1* expression is confined to the gametophore apical cell indicates conservation of the mechanism controlling the SAM despite the difference in the number of stem cells present within the SAM. This study suggests a deep homology among plant pluripotent stem cells in land plants. Acquiring the ability to amplify the stem cells might have been crucial for the evolution of angiosperm SAM. Considering that *LOG* is expressed in multiple cells at the top of the SAM and promotes stem cell activity in angiosperms, increasing the size of the *LOG* expressing region might have been critical for the evolution of the SAM. In angiosperms, cytokinins activate the expression of *WUSCHEL* (*WUS*), a master regulator of stem cell identity within the SAM (46). Specifically, *WUS* regulates the number of stem cells in the SAM in angiosperms (47). However, recent studies suggest that the function of WUS in the SAM has evolved after the divergence of seed plant lineages (17,48–50). Therefore, acquisition of novel factors promoting stem cell identity, including WUS, might also have been necessary for the evolution of a SAM containing multiple stem cells.

**Fig. 5.**
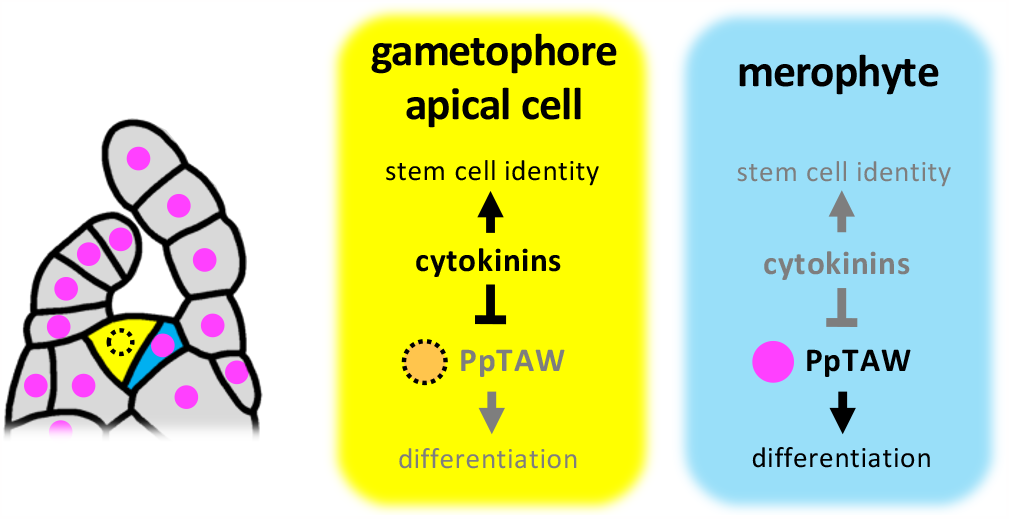
Model for the establishment of asymmetric cell fates of the gametophore apical cell and the merophyte. Cytokinin levels are elevated in the gametophore apical cell where it promotes pluripotent stem cell identity and represses PpTAW protein accumulation. On the other hand, cytokinin levels decrease in the merophyte after cell division, allowing accumulation of PpTAW. PpTAW promotes merophyte identity and facilitates merophyte differentiation. Therefore, the gametophore apical cellspecific accumulation of cytokinins and suppression of PpTAW accumulation by cytokinins are essential for establishing the asymmetry after the cell division of the gametophore apical cell.

## Acknowledgments

We thank Dr. Mitsuyasu Hasebe (National Institute for Basic Biology, Japan) for providing pCit-aphIV, pPIG1b:NGGII, pPGX8, pTN186, pTN182, p35S-loxP-Zeo, and p35S-loxP-BSD vectors. We also thank Dr. Maya Bar (Agricultural Research Organization, Israel) for providing theTCSv2:3xVENUS vector.

## Author contributions

Conceptualization: JK, YHa ; Methodology: YHi, SN ; Investigation: YHa, JO ; Visualization: YHa, JO ; Funding acquisition: JK, SN, YHa ; Project administration: JK ; Supervision: JK ; Writing – original draft: JK, YHa ; Writing – review & editing: JK, YHa

## Data and materials availability

All materials used in the analysis are available upon request subject to a materials transfer agreements. All data are available in the main text or the supplementary materials.

