## Supplementary Materials for "Cytokinins specify pluripotent stem cell identity in the moss *Physcomitrium patens*"

**The PDF file includes:**

Materials and Methods
Figs. S1 to S7
Tables S1

**Other Supplementary Materials for this manuscript include the following:**

Movies S1 to S4

**Materials and Methods**

Plant materials and culture conditions

The Gransden Wood strain (1962) of *Physcomitrium patens* was used as wild type (51). *P. patens* plants were cultured in BCDAT medium, under continuous light at 25℃, as described by Nishiyama et al. 2000, and then transferred to media for analysis (52).

For the observation of plant colonies of the *PpTAW*s multiple mutants, *PpTAW:Citrine* reporter lines, *proPpTAW2:NGG* reporter lines, and *TCSv2:GUS* reporter lines, small amounts of protonemal tissues were cultured between two cellophane membranes (PL#300, Futamura Chemical Co., Ltd.) on BCD medium from 2 to 3 weeks. For the observation of gametophores of the *PpTAW* quadruple mutants, plants were cultured on BCDAT medium for 1 month. For live imaging of *PpTAW2:Citrine;LTI6b:RFP* lines, small amounts of protonemal tissues were placed on thin BCD medium covering the bottom of a glass dish (D11140H, Matsunami Glass Ind., Ltd.) and then covered with a cellophane membrane so that plants grow in the narrow space between the glass bottom and the cellophane membrane. Gametophore production was usually observed from day 10 to 18, and gametophores in optimal developmental stage were selected and used for live imaging.

For the analysis of *XVE:PpTAW2* plants, the plants were grown on the BCD medium containing appropriate concentration of β-estradiol (Fujifilm Wako Pure Chemical Corp.) or a same amount of solvent (DMSO) as a control. Growth conditions were modified depending on the developmental stages to be analyzed. For the observation of plant growth phenotypes, plants were cultured for 3 weeks between two cellophane membranes. For the regeneration assay from detached leaves, gametophores grown on BCDAT medium for 3 weeks were collected and soaked in BCD liquid medium containing the appropriate concentration of β-estradiol or the same amount of solvent (DMSO) for 24 hours. Soaked gametophores were thoroughly washed with sterile water, and the five youngest leaves were cut from each gametophore. Detached leaves were pooled and mixed. Ten leaves were randomly selected and placed on BCDAT medium. Cellophane membranes were placed on top of the leaves and they were cultured for 3 days.

For the observation of PpTAW2-Citrine localization in *XVE:PpTAW2-Citrine* plants, the plants were cultured for 3 weeks between two cellophane membranes, and then cultured in sterilized water containing 100 nM β-estradiol or the same amount of solvent (DMSO) for 24 hours.

For analyses of the phenotypes of *XVE:PpTAW2-SRDX* plants, firstly young gametophores were produced by culture on plain BCD medium plates for 3 weeks, followed by addition of 20 mL of sterilized water containing 100 nM β-estradiol, or the same amount of solvent, and an additional 4 weeks of culture. For the observation of merophyte development, small amounts of protonemal tissues were cultured between two cellophane membranes on BCD medium for 3 weeks and then transferred to BCD medium plates containing appropriate concentrations of β-estradiol or the same amount of solvent (DMSO). The plants were further cultured for 1 week.

Phylogenetic analysis

BlastP searches were performed on Phytozome (https://phytozome.jgi.doe.gov/pz/portal.html) using default parameter settings to look for ALOG family proteins and LOG proteins in *Oryza sativa*, *Arabidopsis thaliana*, *Ceratopteris richardii*, *Selaginella moellendorffii*, *Marchantia polymorpha*, *Physcomitrium patens*, and *Chara braunii*. For the ALOG family proteins, the amino acid sequence of TAW1 from Oryza sativa was used as query in the BlastP search. Amino acid sequences found in BlastP search were aligned using Clustal Omega (https://www.ebi.ac.uk/Tools/msa/clustalo/), and the ALOG domain region without gaps were extracted using the Geneious software to get the multiple sequences alignmentsa for the phylogenetic analysis. For the LOG proteins, the amino acid sequence of LOG from Oryza sativa was used as a query in the BlastP search. Amino acid sequences found in the BlastP search were aligned with Clustal Omega, and the lysine decarboxylase domain region without gaps were extracted using the Geneious software. The phylogenetic analysis was performed on PhyML (http://www.atgc-montpellier.fr/phyml/), based on the maximum likelihood method with 1000 times bootstrap.

Vector construction

The pCit-aphIV vector (gift from Mitsuyasu Hasebe, NIBB) was used to make *PpTAW:Citrine* knock-in constructs for visualization of PpTAW protein localizations. Both DNA fragments around 1.2 kbp upstream and downstream of the stop codon of each *PpTAW* were amplified by PCR. The DNA fragment of the upstream region was cloned into the EcoRV site just before the Citrine coding region in the pCit-aphIV vector so that the reading frame of *PpTAW* is in frame with the Citrine's using a linker sequence (GGAGGAGGATCA). The DNA fragment of the downstream region was cloned into the SmaI site just after the selective marker gene cassette of the pCit-aphIV.

The pPIG1b:NGGII vector (accession number AB537478) was used to make the *proPpTAW2:NGG* and *TCSv2:GUS* constructs (53). For the *proPpTAW2:NGG* construct, DNA fragments including 3.5 kbp around the promoter region of *PpTAW2* were amplified by PCR and cloned into the SmaI site of the pPIG1b:NGGII vector using the SLiCE method (54). For the *TCSv2:GUS* construct, DNA fragments containing the *TCSv2* promoter were amplified using the TCSv2:3xVENUS vector as a template and cloned into the SmaI site of pPIG1b:NGGII (33).

pTN186 (Addgene plasmid #34890), pTN182 (Addgene plasmid #34888), p35S-loxP-Zeo (AB540628), and p35S-loxP-BSD (AB537973) were used to make constructs for the disruption of *PpTAW2*, *PpTAW3*, *PpTAW4*, and *PpTAW1* respectively (13,55). DNA fragments 1.2 kbp upstream and downstream of the coding region of each *PpTAW*, were amplified by PCR. The DNA fragment of the upstream region was cloned into the EcoRV site before the selective marker gene cassette in the vector. The DNA fragment of the downstream region was cloned into the SmaI site after the selective marker gene cassette in the vector.

The pPGX8 vector (AB537482) was used to make *XVE:PpTAW2*, *XVE:PpTAW2-Citrine*, and *XVE:PpTAW2-SRDX* constructs (41). A DNA fragments containing the coding sequence of *PpTAW2* or *PpTAW2-Citrine* were amplified by PCR using genomic DNA of *P. patens* or PpTAW2:Citrine vector as template, respectively. Amplified DNA fragments were subcloned into pENTR/D-TOPO vector (Invitrogen). The sequence of *PpTAW2* or *PpTAW2-Citrine* were transferred to pPGX8 by LR reaction to generate *XVE:PpTAW2* or *XVE:PpTAW2-Citrine* construct. For the construction of *XVE:PpTAW2-SRDX*, the SRDX sequence (CTGGATCTGGATCTGGAACTGCGCCTGGGCTTTGCG) was introduced just before the stop codon of *PpTAW2* subcloned in pENTR/D-TOPO, by site-directed mutagenesis using inverse PCR (40). The sequence of *PpTAW2-SRDX* in pENTR/D-TOPO was transferred to pPGX8 by LR reaction to generate the *XVE:PpTAW2-SRDX* constructs. The list of primers used for vector construction is shown in Table S1.

Transformation and genotyping

Six to seven days-old protonema tissue cultured on BCDAT medium overlaid with cellophane was used for the transformation. Polyethylene glycol mediated transformation was performed as described in Nishiyama et al., 2000 (52). Plasmids used for the transformation were extracted using the Fast gene plasmid mini kit (Nippon Genetics Co., Ltd.). For introduction of *PpTAW:Citrine*, *proPpTAW2:NGG*, *TCSv2:GUS*, *XVE:PpTAW2*, *XVE:PpTAW2-Citrine*, and *XVE:PpTAW2-SRDX* constructs, the plasmids were digested by the appropriate restriction enzymes to isolate the DNA fragments to be integrated into the genome by homologous recombination. For introduction of *PpTAW* genes disruption constructs, DNA fragments to be integrated into the genome by homologous recombination were amplified by PCR using the plasmids as template. 15 ~ 20 µg of plasmid purified by ethanol precipitation were used for the transformation. After selection on the medium containing antibiotics, genomic DNA was extracted from each regenerated plant colony and PCR amplification of the regions inside and outside of the introduced constructs was performed to confirm the integration of the construct in the genome (fig. S6 and S7). The PCR products were checked by agarose gel electrophoresis and lanes showing bands of appropriate length were selected. For the introduction of the *LTI6b:RFP* construct, regenerated plants showing RFP signal at the plasma membrane were manually selected under the fluorescence stereo microscope (M165FC, Leica). The list of primers used for genotyping is shown in Table S1. At least three independent knock-in lines were analyzed for each *PpTAW* gene, and lines showing representative expression patterns were selected for further analysis. Three independent *TCSv2:GUS* marker lines showing similar GUS expression patterns were selected from 9 lines and used to analyze the spatial localization of GUS expression. Three independent *proPpLOG1:GUS* lines showing similar expression patterns of GUS were selected from 9 lines and analyzed.

Histochemical GUS activity assay

GUS staining was conducted following Aoyama et al., 2012 with minor modifications (13). Plant tissues were fixed with fixation solution [0.2% (w/v) MES (pH 5.6), 0.3% (v/v) Formalin, 0.3 M Mannitol] at room temperature for 10 minutes. After washing with 50 mM $NaH_2PO_4$ (pH 7.0), the fixed tissues were vacuum-infiltrated with a substrate solution [50 mM $NaH_2PO_4$ (pH 7.0), 0.5 mM 5-bromo-4-chloro-3-indolyl β-D-glucuronide (X-Gluc), 0.5 mM $K_3Fe(CN)_6$, 0.5 mM $K_4Fe(CN)_6$, and 0.05% (v/v) Triton X-100] for 30 minutes, and stained at 37°C. After the staining, the tissues were fixed with 5% (v/v) formalin for 10 minutes and then soaked in 5% (v/v) acetic acid for 10 minutes. The stained and fixed tissues were dehydrated with ethanol series. The tissues were cleared by incubation in chloral hydrate solution [66% (w/w) chloral hydrate, 8% (w/w) glycerol] at 4°C overnight before imaging.

Plant embedding and sectioning

GUS-stained plant tissues (dehydrated with ethanol series) were embedded in Technovit 7100 resin (Heraeus Kulzer) following the manufacturer's instructions. Embedded samples were sectioned on a rotary microtome with 7 µm thickness. The obtained sections were treated with neutral red dyes as a counterstain. Multi-Mount 480 solution (Matsunami Glass Ind., Ltd.) was used as mounting agents on the slides.

Microscopy

PpTAW:Citrine fluorescence was observed using a confocal scanning microscope (LSM880, Zeiss). Plan-Apochromat 20x/0.80 M27 objective or LD LCI Plan-Apochromat 40x/1.2 Imm Korr DIC M27 water objective were used. Citrine was excited at 514 nm, whereas RFP or cell walls stained with 50 mg/mL propidium iodide (PI) were excited at 543 nm. Morphology of

*XVE:PpTAW2* plants, *pptaw*s loss-of-function mutants, and *XVE:PpTAW2-SRDX* plants were observed using a stereo microscope (M165FC, Leica) or the confocal scanning microscope (LSM880, Zeiss). For the confocal microscope observations, cell walls stained by PI (50 mg/mL) and chloroplasts' autofluorescence were excited at 543 nm and 633 nm, respectively. The 3D reconstruction of z-stack images was done with the ZEN black software (Zeiss). GUS-stained samples were observed with a light microscope (BX51, Olympus) equipped with an Olympus DP71. UPlanFl 40× objective was used.

<u>Quantitative RT-PCR</u>

Collected plant samples were frozen in liquid nitrogen and crushed to a fine powder using a multi beads homogenizer (Yasui Kikai). Total RNA was extracted from the tissue powder using the NucleoSpin RNA Plant kit (Macherey-Nagel) following the manufacture's instructions. cDNA was synthesized by using the SuperScript VILO cDNA Synthesis Kit (Invitrogen). Real time PCR was performed with KOD SYBR (TOYOBO) on LightCycler480II (Roche). *PpEF1α* (Pp3c2_10310) or *PpACT5* (Pp3c10_17070) were used as reference genes. The list of primers used for qRT-PCR is shown in Table S1.

<u>Data analyses and statistics</u>

Quantitative analysis of images was performed using ImageJ software and statistical analyses were conducted in Excel or R software. For the quantification of the PpTAW2:Citrine signal in the nuclei, nuclei regions were extracted using the Analyze Particle function, after conversion to grayscale. Mean gray value in the nuclei regions were measured. For quantification of the pigmentation in rhizoid, rhizoid cell regions and outside regions surrounding rhizoid cell were manually selected and differences of mean gray value between cell regions and outside regions were calculated. The comparison of the two groups were conducted on Excel and p-values were calculated by student's *t*-test. The comparison of more than two groups were conducted on R and p-values were calculated by performing the HSD (Honestly Significant Differences) test using the multcomp package. All experiments were repeated at least three times.

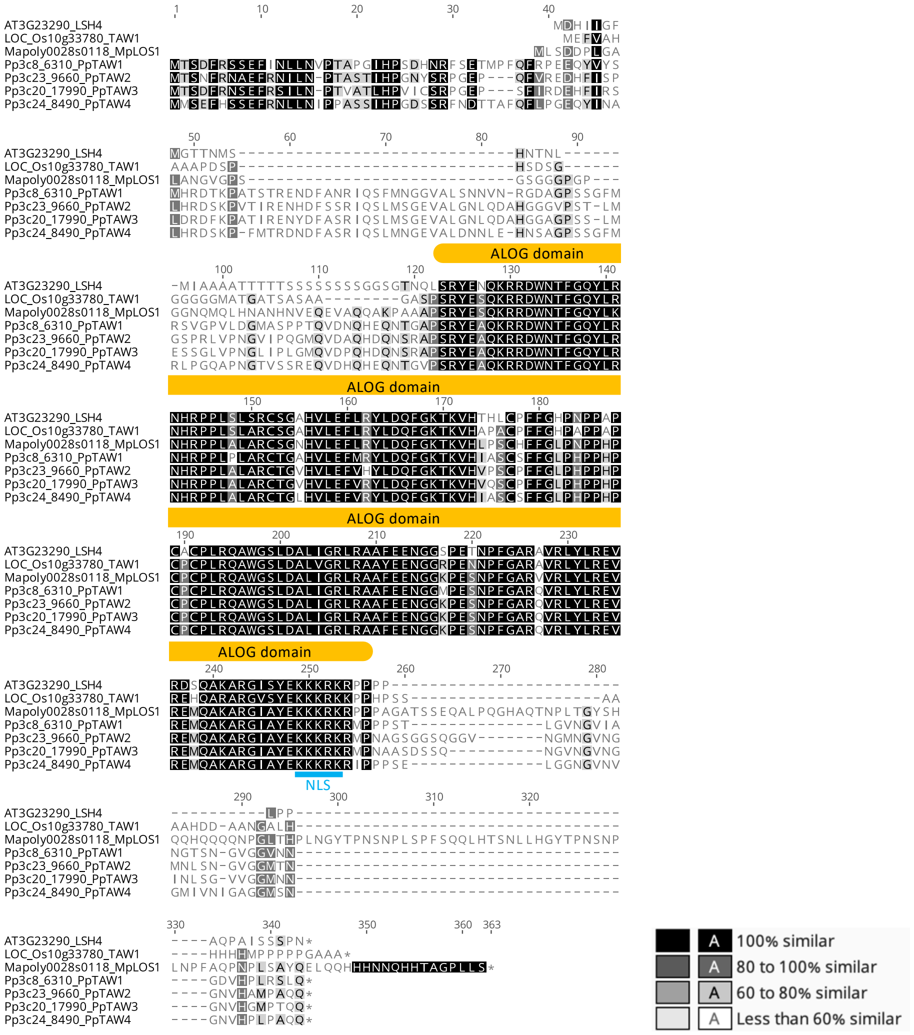

**Fig. S1. Conservation of amino acid sequences in ALOG proteins.**

Alignment of amino acid sequences of ALOG proteins. Species shown in the alignment are *Physcomitrium patens* (Pp), *Marchantia polymorpha* (Mapoly), *Arabidopsis thaliana* (AT), and *Oryza sativa* (Os). Each amino acid letters are highlighted by black colors based on similarity. Region of the ALOG domain is indicated by orange bars. Position of the nuclear localization signal (NLS) is shown by a cyan bar.

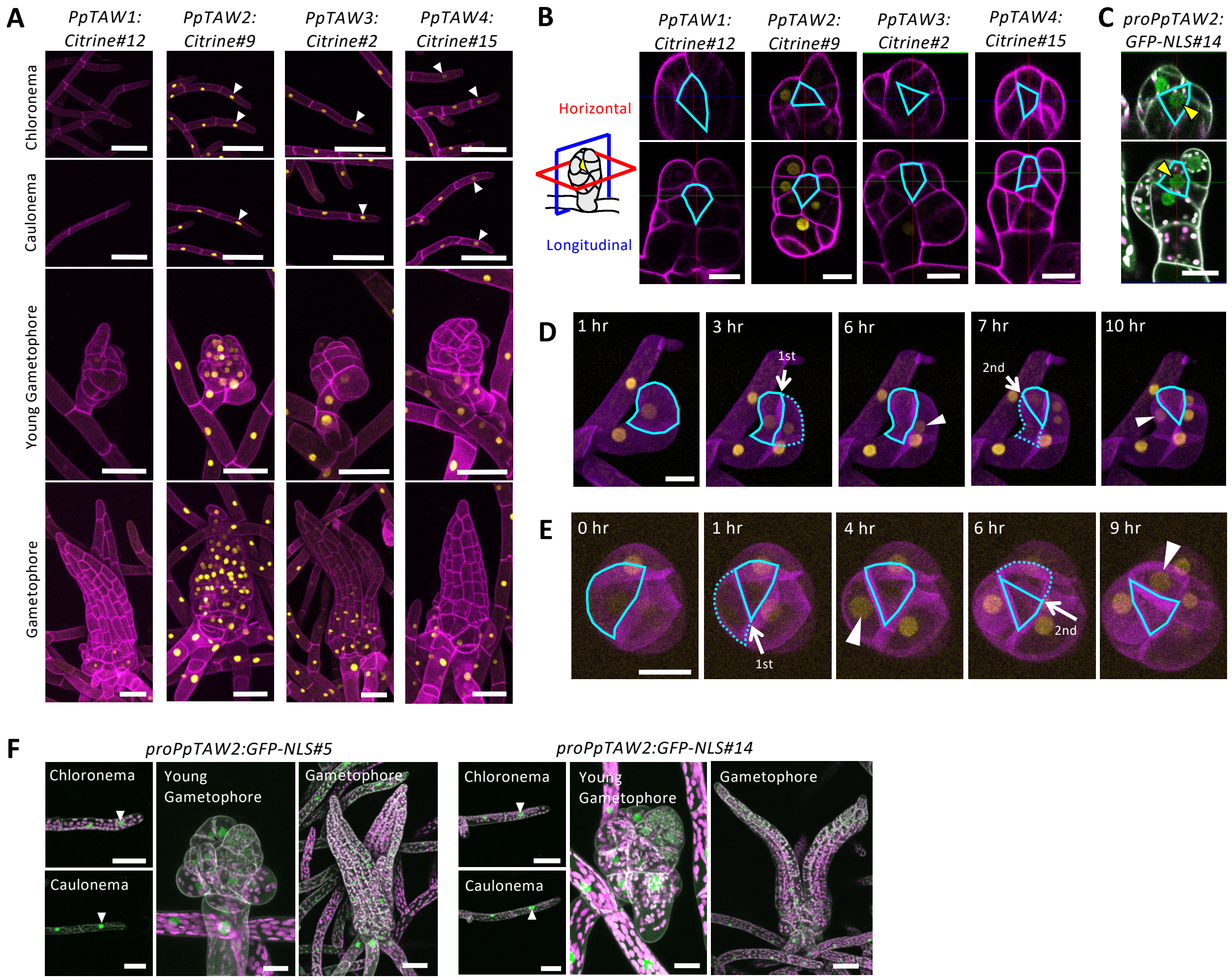

**Fig. S2. Localization of PpTAW proteins and promoter activity of *PpTAW2* in other lines.**

(**A**) Localizations of PpTAW:Citrine fluorescence (yellow) in protonema, young gametophore and gametophore. PpTAW:Citrine fluorescence is visible in the chloronemal and caulonemal apical cells (white arrowheads). Cell walls were stained with propidium iodide (magenta). (**B**) Localizations of PpTAW:Citrine fluorescence (yellow) in the shoot apical meristem (SAM). The top panels show horizontal (red square) views, and the bottom panels show longitudinal (blue square) views. Cell walls are stained with propidium iodide (magenta). The outlines of the gametophore apical cell is marked with cyan lines. (**C**) Promoter activity of *PpTAW2*. GFP fluorescence (green) driven by the *PpTAW2* promoter is shown. Cell walls are stained with propidium iodide (white). The magenta color represents the autofluorescence of chloroplasts. Yellow arrowheads indicate the GFP fluorescence in the gametophore apical cell. (**D** and **E**) Time-lapse imaging of PpTAW2:Citrine (yellow) localization during the division of gametophore apical cells in an initiating gametophore (D) and a growing young gametophore (E). LTI6b:RFP (magenta) was simultaneously imaged for the visualization of cell outlines. Solid and dashed cyan lines show outlines of the gametophore apical cell and newly formed merophyte, respectively. White arrows indicate the cell division planes formed during the observation time. White arrowheads indicate the PpTAW2:Citrine signal that appeared in the merophyte after the division of the gametophore apical cell. No PpTAW2:Citrine signal is observed in the gametophore apical cell. (**F**) Promoter activity of PpTAW2 (green) during developmental processes from protonema to gametophores. Cell walls were stained with propidium iodide (white). Magenta color shows autofluorescence of chloroplasts. White arrowheads represent promoter activity in the chloromena and caulonema stem cell. Scale bars, 50 µm (A, F, Chloronema, Caulonema, and Gametophore), and 20 µm (B to E, and F, Young Gametophore).

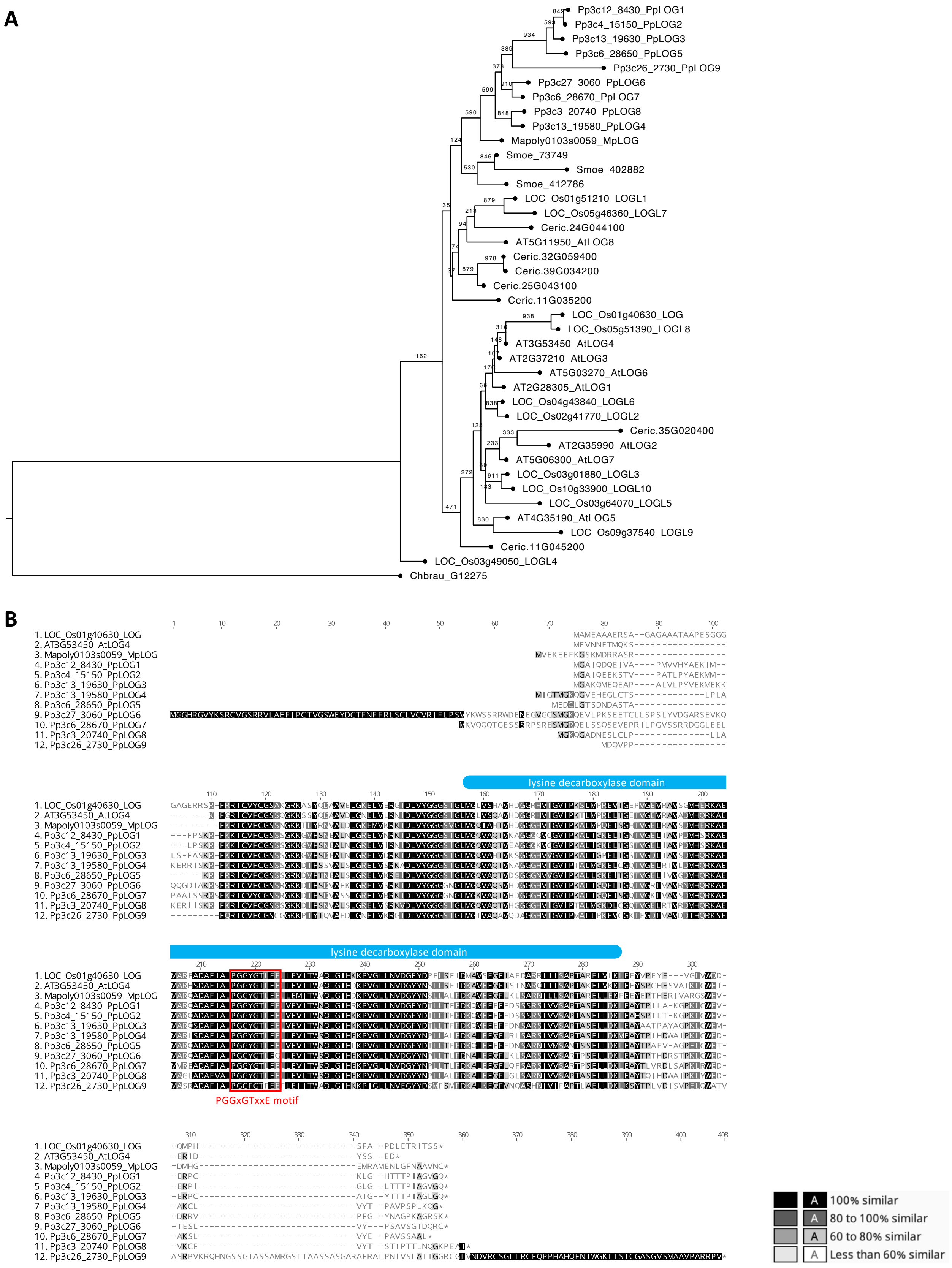
A
Pp3c12_8430_PpLOG1
Pp3c4_15150_PpLOG2
Pp3c13_19630_PpLOG3
Pp3c6_28650_PpLOG5
Pp3c26_2730_PpLOG9
Pp3c27_3060_PpLOG6
Pp3c6_28670_PpLOG7
Pp3c3_20740_PpLOG8
Pp3c13_19580_PpLOG4
Mapoly0103s0059_MpLOG
Smoe_73749
Smoe_402882
Smoe_412786
LOC_Os01g51210_LOGL1
LOC_Os05g46360_LOGL7
Ceric.24G044100
AT5G11950_AtLOG8
Ceric.32G059400
Ceric.39G034200
Ceric.25G043100
Ceric.11G035200
LOC_Os01g40630_LOG
LOC_Os05g51390_LOGL8
AT3G53450_AtLOG4
AT2G37210_AtLOG3
AT5G03270_AtLOG6
AT2G28305_AtLOG1
LOC_Os04g43840_LOGL6
LOC_Os02g41770_LOGL2
Ceric.35G020400
AT2G35990_AtLOG2
AT5G06300_AtLOG7
LOC_Os03g01880_LOGL3
LOC_Os10g33900_LOGL10
LOC_Os03g64070_LOGL5
AT4G35190_AtLOG5
LOC_Os09g37540_LOGL9
Ceric.11G045200
LOC_Os03g49050_LOGL4
Chbrau_G12275
B
1. LOC_Os01g40630_LOG
2. AT3G53450_AtLOG4
3. Mapoly0103s0059_MpLOG
4. Pp3c12_8430_PpLOG1
5. Pp3c4_15150_PpLOG2
6. Pp3c13_19630_PpLOG3
7. Pp3c13_19580_PpLOG4
8. Pp3c6_28650_PpLOG5
9. Pp3c27_3060_PpLOG6
10. Pp3c6_28670_PpLOG7
11. Pp3c3_20740_PpLOG8
12. Pp3c26_2730_PpLOG9
lysine decarboxylase domain
lysine decarboxylase domain
PGGxGTxxE motif
100% similar
80 to 100% similar
60 to 80% similar
Less than 60% similar

**Fig. S3. LOG protein family in land plants.**

(**A**) Phylogenetic tree of LOG proteins in land plants. Species shown in the tree are *Chara braunii* (Chbrau), *Physcomitrium patens* (Pp), *Marchantia polymorpha* (Mapoly), *Selaginella moellendorffii* (Sm), *Ceratopteris richardii* (Ceric), *Arabidopsis thaliana* (AT), and *Oryza sativa* (Os). A LOG protein in *Chara braunii* (Chbrau_G29454) was included as an outgroup. The bootstrap value is indicated beside the branch point. (**B**) Alignment of amino acid sequences of LOG proteins. Each amino acid letters are highlighted by black colors based on similarity. Region of the lysine decarboxylase domain is indicated by blue bars. Position of the conserved PGGxGTxxE motif is shown by a red box.

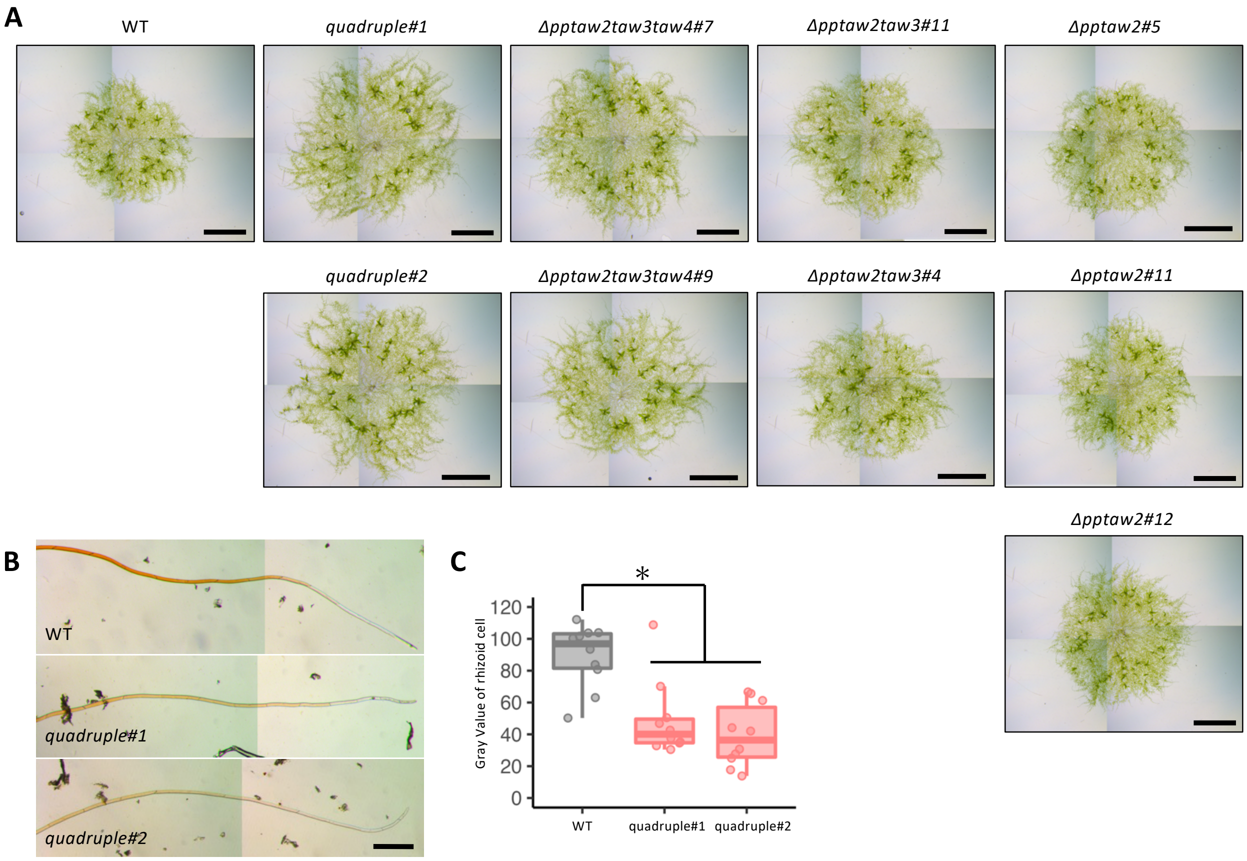

**Fig. S4. Phenotype of *PpTAW*s loss-of-function mutants.**

(**A**) 23 days plant colony of WT and the *PpTAW*s loss-of-function mutants. (**B**) Pigmentation of rhizoids in the WT and quadruple mutants. (**C**) Gray value of the 10th rhizoid cell from tip in the in the WT and quadruple mutants. Statistical significance was assessed by student's t-test ($n = 10$, $p < 0.05$). Scale bars, 4 mm (A), and 200 µm (B).

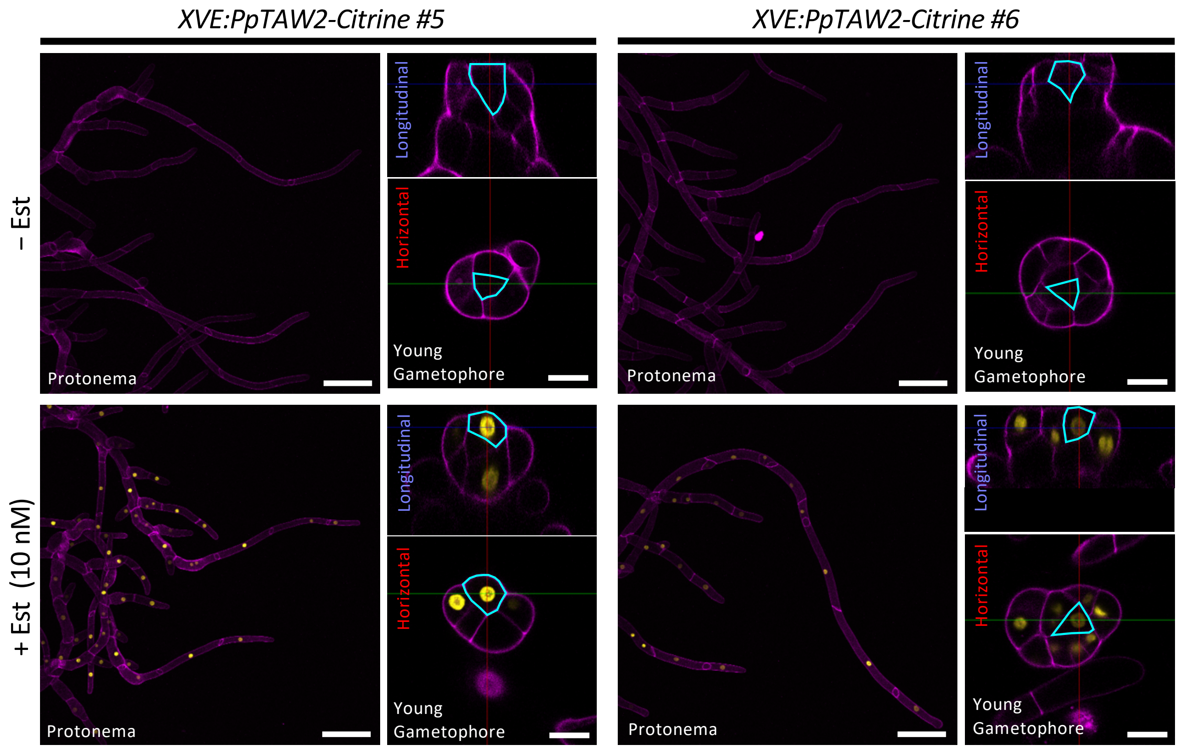

**Fig. S5. Ectopic localization of PpTAW2 protein.**

The β-estradiol inducible system of PpTAW2-Citrine. β-estradiol treatment (10 nM, 24 hours) induced the expression of the PpTAW2-Citrine (yellow) localized to nuclei in both protonema and young gametophores including the gametophore apical cell indicated by cyan lines. No signal was detected in the control. Cell walls were stained with propidium iodide staining (magenta). Scale bars, 100 μm in protonema images, and 20 μm in young gametophore images.

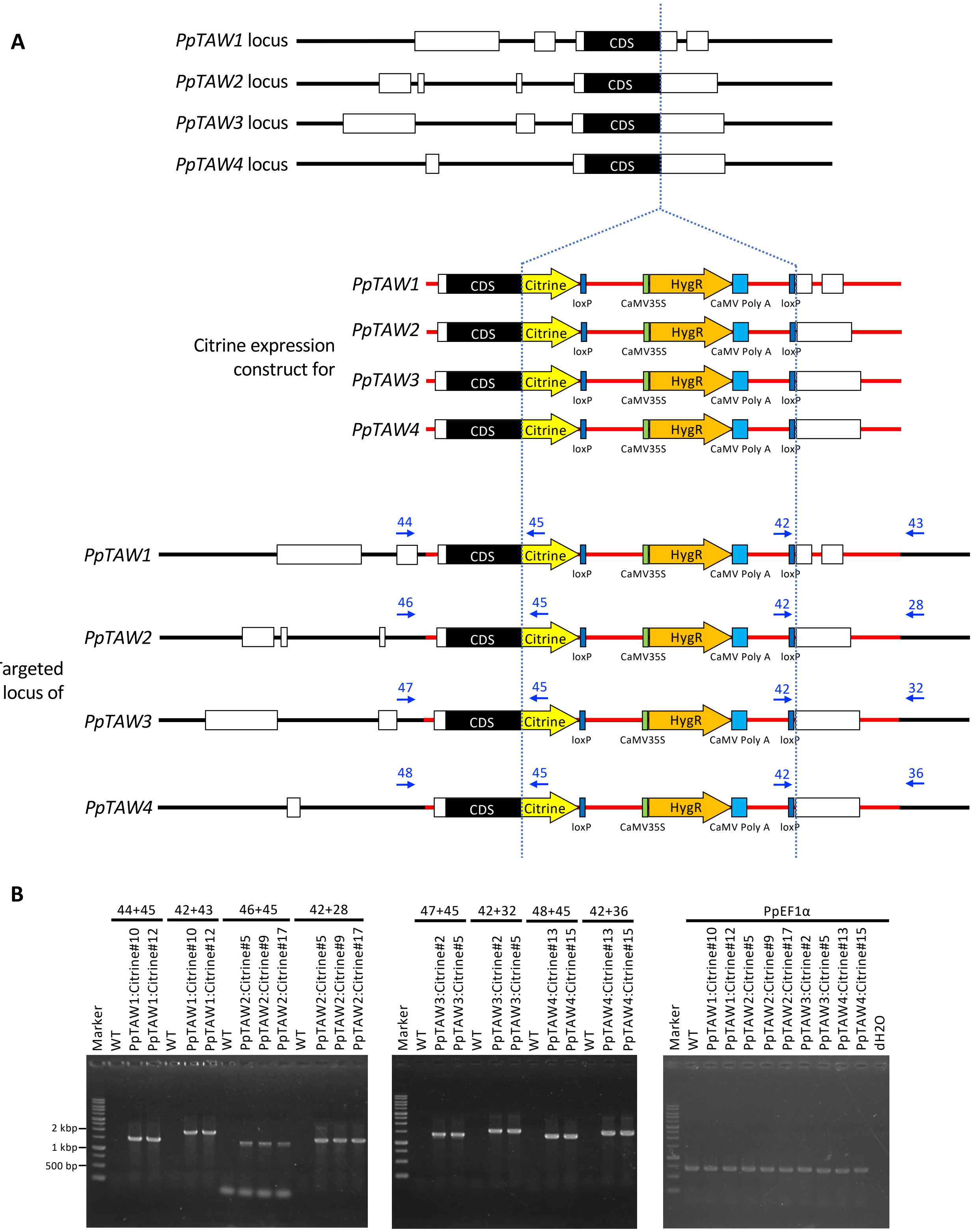

**Fig. S6. Construction for visualization of PpTAW1-4 localization by Knocking in Citrine.** (**A**) Schematic diagrams of *PpTAW1-4* loci and targeting of the constructs. White and black boxes indicate exons and protein- coding sequences (CDS), respectively. Blue arrows with a number show the position of primers used in genotyping. The numbers are corresponding to the primer numbers in Table S1. (**B**) Examination of PCR products for the genotyping by the agarose gel electrophoresis. Numbers of the top show primer pairs used in the PCR. The numbers are corresponding to the primer numbers in Table S1. Genomic region encoding *PpEF1α* was amplified simultaneously as a positive control of the PCR (primer #53 and #54 were used).

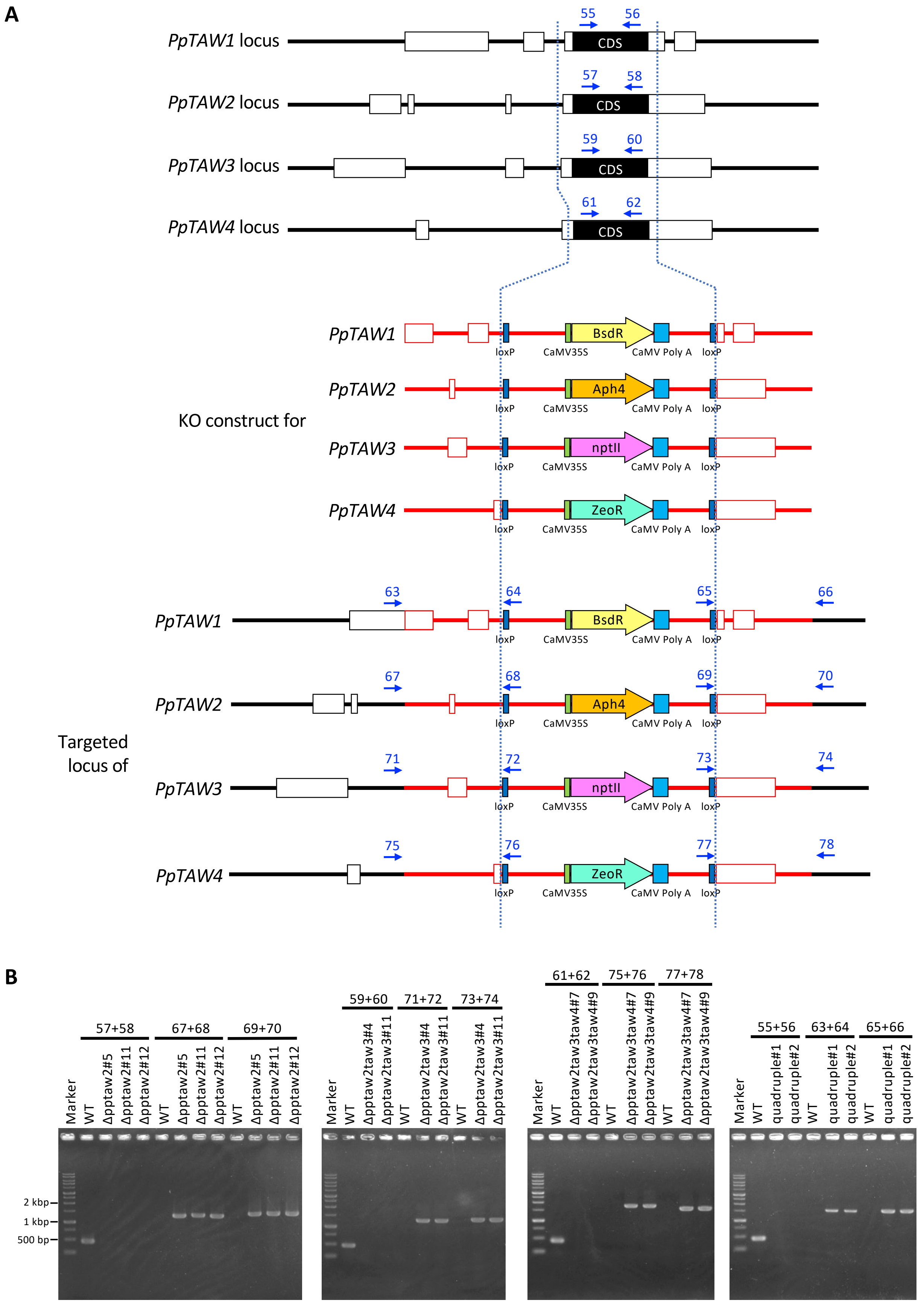
A
PpTAW1 locus
PpTAW2 locus
PpTAW3 locus
PpTAW4 locus
CDS
55
56
57
58
59
60
61
62
KO construct for
PpTAW1
PpTAW2
PpTAW3
PpTAW4
BsdR
Aph4
nptII
ZeoR
loxP
CaMV35S
CaMV Poly A
Targeted locus of
B
57+58
67+68
69+70
Marker
WT
Δpptaw2#5
Δpptaw2#11
Δpptaw2#12
2 kbp
1 kbp
500 bp
59+60
71+72
73+74
Δpptaw2taw3#4
Δpptaw2taw3#11
61+62
75+76
77+78
Δpptaw2taw3taw4#7
Δpptaw2taw3taw4#9
55+56
63+64
65+66
quadruple#1
quadruple#2

**Fig. S7. Construction for *PpTAW1-4* KO lines.**

(**A**) Schematic diagrams of *PpTAW1-4* loci and targeting of the constructs. White and black boxes indicate exons and protein- coding sequences (CDS), respectively. Blue arrows with a number show the position of primers used in genotyping. The numbers are corresponding to the primer numbers in Table S1. (**B**) Examination of PCR products for the genotyping by the agarose gel electrophoresis. Numbers of the top show primer pairs used in the PCR. The numbers are corresponding to the primer numbers in Table S1. Firstly, *Δpptaw2* single mutants were obtained and *Δpptaw2Δpptaw3* double mutants were generated from *Δpptaw2#5. Δpptaw2Δpptaw3Δpptaw4* triple mutants were generated from *Δpptaw2Δpptaw3#11.* Finally, quadruple mutants were generated from *Δpptaw2Δpptaw3Δpptaw4#9*.

### Table S1. Primers used in this study

| # | primer name | sequence (5'→3') | description |
|---|---|---|---|
| 1 | HRCit5'TAW1clo_F | ATCGTATGCATTTATATAACCATT | for construction of PpTAW1:Citrine |
| 2 | HRCit5'TAW1clo_R | TGATCCTCCTCCCTGCAGTGACCGGAGAG | for construction of PpTAW1:Citrine |
| 3 | HRCit3'TAW1clo_F | TAACAAAGTTTTGACCTCCGAA | for construction of PpTAW1:Citrine |
| 4 | HRCit3'TAW1clo_R | GATATCGATTGAAGAGGCGGT | for construction of PpTAW1:Citrine |
| 5 | HRCit5'TAW2clo_F | ATCACATTCACATTTGTAGTGC | for construction of PpTAW2:Citrine |
| 6 | HRCit5'TAW2clo_R | TGATCCTCCTCCTTGCTGCGCCGGCATGG | for construction of PpTAW2:Citrine |
| 7 | HRCit3'TAW2clo_F | GCGAACTGTTTGAACAGAC | for construction of PpTAW2:Citrine |
| 8 | HRCit3'TAW2clo_R | GATATCAATTCTTTCCGCTCA | for construction of PpTAW2:Citrine |
| 9 | HRCit5'TAW3clo_F | ATCACATTCGCGGAGATA | for construction of PpTAW3:Citrine |
| 10 | HRCit5'TAW3clo_R | TGATCCTCCTCCCTGCTGCGTCGGCATGC | for construction of PpTAW3:Citrine |
| 11 | HRCit3'TAW3clo_F | GCGAACTGTTTGAACATACG | for construction of PpTAW3:Citrine |
| 12 | HRCit3'TAW3clo_R | GATATCGTCGATTGATGCGAA | for construction of PpTAW3:Citrine |
| 13 | HRCit5'TAW4clo_F | ATCGATATGTAGCAAACGAC | for construction of PpTAW4:Citrine |
| 14 | HRCit5'TAW4clo_R | TGATCCTCCTCCCTGCTGTGCTGGGAGAG | for construction of PpTAW4:Citrine |
| 15 | HRCit3'TAW4clo_F | GTACAATGCTCTGATCTGCG | for construction of PpTAW4:Citrine |
| 16 | HRCit3'TAW4clo_R | GATATCCAGAAGGCAGAAGCA | for construction of PpTAW4:Citrine |
| 17 | proPpTAW2_clo_F | AGGATCCCCCCCCCCTGGAAGGGCACAAGGAACTG | cloning PpTAW2 promoter |
| 18 | proPpTAW2_clo_R | TGGATCAGCCATCCCCCAAAGATACTGGCTCAGCGA | cloning PpTAW2 promoter |
| 19 | M13F | TGTAAAACGACGGCCAGT | amplification of TCSv2 |
| 20 | TCSv2_in_3xVENUS_reverse | CGATTTCGAACCCGGGGTAC | amplification of TCSv2 |
| 21 | PpTAW1_5'UTR_CloF | GTCTTGCCGATGCTGGTTTG | for construction of PpTAW1 KO |
| 22 | PpTAW1_5'UTR_CloR | GAAAGAAACACTGCGCACCA | for construction of PpTAW1 KO |
| 23 | PpTAW1_3'UTR_CloF | GCTTCCATGCGCAAAGGAAA | for construction of PpTAW1 KO |
| 24 | PpTAW1_3'UTR_CloR | CTTCTTCACGGGCAGGACTC | for construction of PpTAW1 KO |
| 25 | PpTAW2_5'UTR_CloF | TTTAGCCCTGCACTCGTTGT | for construction of PpTAW2 KO |
| 26 | PpTAW2_5'UTR_CloR | GCAGGGGGTATACGATGCTC | for construction of PpTAW2 KO |
| 27 | PpTAW2_3'UTR_CloF | GTCACAGTCCCGTCACAGTT | for construction of PpTAW2 KO |
| 28 | PpTAW2_3'UTR_CloR | TGTCTTCTGCTTGCCTTGCT | for construction of PpTAW2 KO |
| 29 | PpTAW3_5'UTR_CloF | TGCGTTAGCTTGGAACCTGT | for construction of PpTAW3 KO |
| 30 | PpTAW3_5'UTR_CloR | GCAGGAAGCCGAGCTCATAA | for construction of PpTAW3 KO |
| 31 | PpTAW3_3'UTR_CloF | CACAAAGACGACGAGGACGA | for construction of PpTAW3 KO |
| 32 | PpTAW3_3'UTR_CloR | GCGTATGAGTATGGCAGGCT | for construction of PpTAW3 KO |
| 33 | PpTAW4_5'UTR_CloF | ACTGTTTCAGCCCTCACGAG | for construction of PpTAW4 KO |
| 34 | PpTAW4_5'UTR_CloR | CGTAGCGAAGAAGCACTTGC | for construction of PpTAW4 KO |
| 35 | PpTAW4_3'UTR_CloF | TCTGATCTGCGAAACAGCGA | for construction of PpTAW4 KO |
| 36 | PpTAW4_3'UTR_CloR | GCTCTGCAGACTTCTTCGGA | for construction of PpTAW4 KO |
| 37 | topoPpTAW2clo_F | CACCATGACGAGTAACTTTCGCA | cloning PpTAW2 CDS |
| 38 | topoPpTAW2clo_R | TCATTGCTGCGCCGGCA | cloning PpTAW2 CDS |
| 39 | pDONR_PpTAW2_SRDXintr_F | CTGCGCCTGGGCTTTGCGTGAAAGGGTGGGCGCGC | for SRDX introduction |
| 40 | pDONR_PpTAW2_SRDXintr_R | TTCCAGATCCAGATCCAGTTGCTGCGCCGGCATGG | for SRDX introduction |
| 41 | Citrine_clo_R | TTACTTGTACAGCTCGTCCATG | amplification of PpTAW2:Citrine |
| 42 | HRCitT1geno3_F | AGTTATCCCTCACACCGGTG | targeting check for PpTAW:Citrine lines |
| 43 | HRCitT1geno3_R | CGGTGAGTTGGTCTGACACA | targeting check for PpTAW:Citrine lines |
| 44 | HRCitT1geno5_F | CCCCGGTTCTGATCGTGTTT | targeting check for PpTAW:Citrine lines |
| 45 | HRCitT1geno5_R | TGAACTTGTGGCCGTTTACG | targeting check for PpTAW:Citrine lines |
| 46 | CRIT2seq.ck_F_2 | GAGCATCGTATACCCCCTGC | targeting check for PpTAW:Citrine lines |
| 47 | HRCitT3geno5_F | TTATGAGCTCGGCTTCCTGC | targeting check for PpTAW:Citrine lines |
| 48 | CRIT4seq.ck_F_4 | GAGTGCTTGTCTTGGCTTCC | targeting check for PpTAW:Citrine lines |
| 49 | HR_pPGX8_5'geno_F | TGCAACATTTGGAGTTGGCA | targeting check for promoter:NGG lines |
| 50 | HR_pPIG1b_NGGII_5'geno_R | GGGGGATCCACTAGTTCTAGA | targeting check for promoter:NGG lines |
| 51 | HR_pPGX8_3'geno_F | ATAATCCGCATAAGCCCCCG | targeting check for promoter:NGG lines |
| 52 | HR_pPGX8_3'geno_R | TGCATAGTTTAATTAGCAGAGTTGT | targeting check for promoter:NGG lines |
| 53 | semi_qPCR_EF1alpha_F | TGTGGAAGTTCGAGACCGTG | for positive control of PCR |
| 54 | semi_qPCR_EF1alpha_R | GCTTGTCAGACGGCCTTTTG | for positive control of PCR |
| 55 | CRIT1-2nd-alle.seq.ck_F | CATGAGATCAGTCGGACCGG | deletion check for PpTAW1 KO lines |
| 56 | CRIT1-2nd-alle.seq.ck_R | CCTGGCCTTAGCTTGCATCT | deletion check for PpTAW1 KO lines |
| 57 | CRIT2-2nd-alle.seq.ck_F | TCAAGCCGGATACAGTCCCT | deletion check for PpTAW2 KO lines |
| 58 | CRIT2-2nd-alle.seq.ck_R | CTCTCAGGTTTGCCCCCATT | deletion check for PpTAW2 KO lines |
| 59 | CRIT3-2nd-alle.seq.ck_F | AGTGGTGAAGTTGCCCTTGG | deletion check for PpTAW3 KO lines |
| 60 | CRIT3-2nd-alle.seq.ck_R | CAAGTACAGACGCACTTGGC | deletion check for PpTAW3 KO lines |
| 61 | CRIT4-2nd-alle.seq.ck_F | ACGACTTTGCAAGCCGGATA | deletion check for PpTAW4 KO lines |
| 62 | CRIT4-2nd-alle.seq.ck_R | TCAAATGCGGCTCGTAACCT | deletion check for PpTAW4 KO lines |
| 63 | HR_KOT1_5'geno_F | GTGCAGAAGTACCAGCTTGT | targeting check for PpTAW1 KO lines |
| 64 | HR_KOT1_5'geno_R | GGGGGCTGCAGGAATATAACT | targeting check for PpTAW1 KO lines |
| 65 | HR_KOT1_3'geno_F | AGTTATCCCTCACACCGGTG | targeting check for PpTAW1 KO lines |
| 66 | HR_KOT1_3'geno_R | TGCTTGCATACTCGGATTCTCT | targeting check for PpTAW1 KO lines |
| 67 | HR_KOT2_5geno_F | CGTTGCTTAGAAAACCCGCA | targeting check for PpTAW2 KO lines |
| 68 | HR_KOT2_5geno_R | GGGGGCTGCAGGAATATAACT | targeting check for PpTAW2 KO lines |
| 69 | HR_KOT2_3geno_F | AGTTATCCCTCACACCGGTG | targeting check for PpTAW2 KO lines |
| 70 | HR_KOT2_3geno_R | CATAGTTGGCGATGTGTGCG | targeting check for PpTAW2 KO lines |
| 71 | HR_KOT3_pTN182_5'geno_F | TGGCTGTTTCTGTTGGTACCA | targeting check for PpTAW3 KO lines |
| 72 | HR_KOT3_pTN182_5'geno_R | TGACATTTTTGGAGTAGGGGG | targeting check for PpTAW3 KO lines |
| 73 | HR_KOT3_pTN182_3'geno_F | AGTTATCCCTCACACCGGTG | targeting check for PpTAW3 KO lines |
| 74 | HR_KOT3_pTN182_3'geno_R | GTATGCAGATGTACGCGACG | targeting check for PpTAW3 KO lines |
| 75 | HRT4geno5'_F | TTTGTGCTCGTTGACGCTTG | targeting check for PpTAW4 KO lines |
| 76 | HRT4geno3'_R | CAAAAACGACCTCACCAGCC | targeting check for PpTAW4 KO lines |
| 77 | pTN186_SmaI_seq.check.F | AGTTATCCCTCACACCGGTG | targeting check for PpTAW4 KO lines |
| 78 | pZeo_5insert_ck_R | AGCCCTTTGGTCTTCTGAGA | targeting check for PpTAW4 KO lines |
| 79 | HR_pPGX8_5'geno_F | TGCAACATTTGGAGTTGGCA | targeting check for β-estradiol inducible lines |
| 80 | HR_pPGX8_5'geno_R | TGTACAGTACGTCGAGGGGA | targeting check for β-estradiol inducible lines |
| 81 | HR_pPGX8_3'geno_F | ATAATCCGCATAAGCCCCCG | targeting check for β-estradiol inducible lines |
| 82 | HR_pPGX8_3'geno_R | TGCATAGTTTAATTAGCAGAGTTGT | targeting check for β-estradiol inducible lines |
| 83 | qPCRforPpTAW2_No.2_F | TGCACAGGGGTCCATGTGTT | for qRT-PCR of PpTAW2 |
| 84 | qPCRforPpTAW2_No.2_R | GGTGCGGCAACCCAAAGAAA | for qRT-PCR of PpTAW3 |
| 85 | qPCRforPpCKX1_F2 | GGCGGCACTCTATCGAATGC | for qRT-PCR of PpCKX1 |
| 86 | qPCRforPpCKX1_R2 | GTAGGCGTGCATTGTACCACC | for qRT-PCR of PpCKX1 |
| 87 | qPCRforPpACT5_F | GTCAACAGCGGATTCCCAGC | for qRT-PCR of PpACT5 |
| 88 | qPCRforPpACT5_R | ACCCTCTCCGGCCATATTGC | for qRT-PCR of PpACT5 |
| 89 | GUS_pPCR_2_F | TCTACTTTACTGGCTTTGGTCG | for qRT-PCR of GUS |
| 90 | GUS_pPCR_2_R | CGTAAGGGTAATGCGAGGTAC | for qRT-PCR of GUS |
| 91 | qPCRforPpEF1a_No2_F | ACGCGTTGTTGGCTTTCACT | for qRT-PCR of PpEF1α |
| 92 | qPCRforPpEF1a_No2_R | GCACTGGAGTACTTCGGGGT | for qRT-PCR of PpEF1α |

**Movie S1.**

Time-lapse movie of PpTAW2:Citrine (yellow) localization during the division of gametophore apical cells in an initiating gametophore in Fig. 1I. LTI6b:RFP (magenta) was simultaneously imaged for the visualization of cell outlines. A scale bar, 20 µm.

**Movie S2.**

Time-lapse movie of PpTAW2:Citrine (yellow) localization during the division of gametophore apical cells in an initiating gametophore in fig. S2D. LTI6b:RFP (magenta) was simultaneously imaged for the visualization of cell outlines. A scale bar, 20 µm.

**Movie S3.**

Time-lapse movie of PpTAW2:Citrine (yellow) localization during the division of gametophore apical cells in a growing young gametophore in Fig. 1J. LTI6b:RFP (magenta) was simultaneously imaged for the visualization of cell outlines. A scale bar, 20 µm.

**Movie S4.**

Time-lapse movie of PpTAW2:Citrine (yellow) localization during the division of gametophore apical cells in a growing young gametophore in fig. S2E. LTI6b:RFP (magenta) was simultaneously imaged for the visualization of cell outlines. A scale bar, 20 µm.
